# Tyrosine-Derived Polycarbonate Nerve Guidance Tubes Elicit Pro-Regenerative Extracellular Matrix Deposition When Used to Bridge Segmental Nerve Defects in Swine

**DOI:** 10.1101/2020.01.11.902007

**Authors:** JC Burrell, D Bhatnagar, DP Brown, NS Murthy, J Dutton, KD Browne, FA Laimo, Z Ali, JM Rosen, HM Kaplan, J Kohn, DK Cullen

**Author notes:** <u>Corresponding author</u>: D. Kacy Cullen, Ph.D., 105E Hayden Hall/3320 Smith Walk, Dept. of Neurosurgery, University of Pennsylvania, Philadelphia, PA 19104, Fx.

## Abstract

Promising biomaterials should be tested in appropriate large animal models that recapitulate human inflammatory and regenerative responses. Previous studies have shown tyrosine-derived polycarbonates (TyrPC) are versatile biomaterials with a wide range of applications across multiple disciplines. The library of TyrPC has been well studied and consists of thousands of polymer compositions with tunable mechanical characteristics and degradation and resorption rates that are useful for nerve guidance tubes (NGTs). NGTs made of different TyrPCs have been used in segmental nerve defect models in small animals. The current study is an extension of this work and evaluates NGTs made using two different TyrPC compositions in a 1 cm porcine peripheral nerve repair model. We first evaluated a nondegradable TyrPC formulation, demonstrating proof-of-concept chronic regenerative efficacy up to 6 months with similar nerve/muscle electrophysiology and morphometry to the autograft repair control. Next, we characterized the acute regenerative response using a degradable TyrPC formulation. After 2 weeks *in vivo*, TyrPC NGT promoted greater deposition of pro-regenerative extracellular matrix (ECM) constituents (in particular collagen I, collagen III, collagen IV, laminin and fibronectin) compared to commercially available collagen-based NGTs. This corresponded with dense Schwann cell infiltration and axon extension across the lumen. These findings confirmed results reported previously in a mouse model and reveal that TyrPC NGTs were well tolerated in swine and facilitated host axon regeneration and Schwann cell infiltration in the acute phase across segmental defects - likely by eliciting a favorable neurotrophic ECM milieu. This regenerative response ultimately can contribute to functional recovery.

## Introduction

Peripheral nerve injury (PNI) often results in insufficient functional recovery following surgical repair [1]. Current PNI repair strategies remain inadequate due to inherent regenerative challenges that are dependent on the severity and/or gap-length of the nerve injury. Although small defects can be repaired by suturing the nerve stumps directly together, longer defects require a bridging graft to act as a guide and protective encasement for regenerating axons [2]. Nerve transection results in complete axon degeneration distal to the injury, necessitating axonal regeneration over long distances to reinnervate target end-organs. These long distances are compounded by the relatively slow growth of regenerating axons (~1 mm/day), diminishing the likelihood for significant functional recovery [1].

Nerve autografts are the gold standard for segmental nerve repair. However, various issues have been associated with suboptimal outcomes, including limited availability of donor nerves, donor site morbidity, diameter mismatch, fascicular misalignment, and modality mismatch (sensory nerves are usually used to repair motor/mixed nerve defects) [3]. Despite advancements over the last few decades, there has been limited success in developing a suitable replacement for autografts. Several nerve gap repair solutions are commercially available (e.g., Stryker Neuroflex™, Baxter GEM NeuroTube®, Axogen Avance® acellular nerve allograft), however clinical use remains limited primarily to noncritical sensory nerve injuries, with autografts still being used widely for motor and critical sensory nerve repairs [4]. Thus, there is a clinical unmet need for a technology that at least matches the autograft repair in functional recovery.

Nerve guidance tubes (NGTs), such as Neuroflex or NeuroTube, are artificial nerve grafts for segmental nerve repair that can be fabricated from biologic or synthetic polymers [2, 5]. Natural polymers offer a micro-environment that facilitate cell attachment but often have undesirable mechanical properties. Synthetic materials are an attractive alternative source, which in addition to promoting nerve regeneration, can provide more desirable mechanical properties and reduced batch-to-batch variability that may potentially lower the overall cost for commercialization. Previous reports have demonstrated efficacy of NGTs comprised of degradable polymers in large animal models [6]. More recently, braiding of polymer fibers has been introduced to generate porous NGTs with mechanical properties and flexibility suitable for nerve repair [7–9]. In this study, a braided NGT was fabricated using two polymers chosen from a library of tyrosine-derived polycarbonates (TyrPCs), which have previously been shown to facilitate nerve regeneration in rodent models and demonstrated improved functional recovery in small gap injuries [10–12]. While braiding may improve mechanical strength and increase kink-resistance, macropore structures may form allowing for tissue infiltration into the conduit that can hinder nerve regeneration. To address these concerns, TyrPC NGTs were coated with a biocompatible hyaluronic acid (HA)-based hydrogel with sufficient porosity for mass transport across the conduit wall yet prohibiting cellular infiltration into the graft *in vivo* [9, 12].

Although previous rodent studies suggest TyrPC NGTs may be useful for bridging segmental defects following PNI; the advancement of new materials for potential clinical translation requires thorough investigation in appropriate large animal models. Indeed, a major consideration for development of novel, clinically translatable NGTs is the inability of rodent models to fully replicate human segmental nerve defects due to critical differences in biological physiology and the short regenerative distances to distal end-target relative to the longer distances implicated in poor functional recovery in humans [13]. Thus, to evaluate whether a novel biomaterial designed for nerve regeneration has potential for clinical translation, demonstration of regenerative efficacy and evaluation of host responses should be performed using in a suitable large animal pre-clinical model [14]. The transition of a novel biomaterial from evaluation in a small animal model to a large animal model typically follows a two-step process—the critical first step is to investigate acute regeneration in a short (e.g., 1 cm) nerve gap. At this stage, it is necessary to demonstrate safety and tolerability, ensuring that there is no deleterious inflammatory response, and also efficacy in repairing short gap injuries. The next step is to demonstrate efficacy as a repair strategy for a challenging long gap nerve defect. Here, we report the first phase in the evaluation of braided NGTs for peripheral nerve injury in a 1 cm porcine nerve injury model. We first performed a proof-of-concept, 6-month study in a porcine model to evaluate functional recovery using a nondegradable TyrPC, and then compared the pro-regenerative capabilities of degradable TyrPC to commercially-available collagen NGTs at an acute time point.

## Methods

### TyrPC NGT Fabrication

TyrPC polymers are composed of desaminotyrosyl-tyrosine (DT), desaminotyrosy-ltyrosine ethyl ester (DTE), and different amounts of PEG. TyrPC nomenclature follows the pattern Exxyy (*n*k), where xx and yy are percentage mole fractions of DT and PEG respectively, and *n* is the molecular weight of PEG in kDa [15, 16]. E0000 is a nondegradable homopolymer, poly(DTE carbonate), while E1001(1k) is a degradable terpolymer containing 10% DT and 1% PEG (1 kDa). The main difference between E0000 and E1001(1k) is the rate of degradation. At 37 °C and in PBS, the molecular weight decreases to 20% of its starting value in 800 days with E0000 [15, 17] and in 80 days in E1001(1k) (Unpublished). Therefore, for the purposes of this study, E0000 can be regarded as non-degrading over the time course of this 6-month study. In contrast, E1001(1k), which has degradation rate of 9 to 12 months (depending on implant size and shape) is significantly degraded over 6 months.

TyrPC polymers were synthesized as described by Magno et al. [15]. The fabrication of the TyrPC NGTs is a 3-step process consisting of fiber extrusion, braiding of the NGT, and finally the application of a barrier coating, as described by Bhatnagar et al. [12]. The previously published procedure for the fabrication of HA-coated NGTs was followed with some modifications to adjust the NGT diameter to the size required for the swine model [12]. TyrPC polymer powder was melt-extruded into 80–110 μm diameter fibers on a single-screw extruder (Alex James & Associates, Inc., Greenville, SC). The extruded fibers were amorphous, as was the polymer, but the polymer chains were oriented to some degree. The degree of orientation as determined by x-ray diffraction methods was 0.32 on a scale of 0 to 1 [18]. This degree of orientation did not cause any significant shrinkage (reduction in length was less than 1% during the cleaning procedure). These fibers were braided into a tubular NGT (three filaments per yarn; 24 carriers, three twisted fibers/carrier, and traditional 2-over-2 braid) using a tubular braiding machine (ATEX, Technologies Inc., Pinebuff, NC). NGTs were fabricated by braiding the fibers over a 2.3 mm inner diameter Teflon mandrel (Applied Plastics Co., Inc. Norwood, MA) to create NGT walls with a uniform pore size distribution (65 ± 19 μm). Next, NGTs were sonicated in cyclohexane (1x), followed by 0.5% (v/v) TWEEN 20 in DI water (1x), and a final rinse in DI water (5x), and were then vacuum dried overnight at room temperature. The braided NGTs showed no further shrinkage in length or diameter at 37 °C. The cleaned, dried NGTs were subsequently cut with a thermal cutter to 1.2 cm length and the fibers at either end of the NGT were fused to facilitate suturing during implantation. TyrPC NGTs were then treated under UV light for 45 minutes.

### Hyaluronic Acid (HA) Hydrogel Coating and Sterilization

Although braiding may improve mechanical strength and provide kink resistance, infiltration through macropores formed in the conduit wall may disrupt ongoing nerve regeneration [9, 12]. Therefore, braided TyrPC NGTs were coated with a biocompatible hyaluronic acid (HA)-based hydrogel that allows for nutrient transfer across the conduit walls yet prevents infiltration into the graft [9, 12]. Briefly, the outer surface of the braided NGTs was coated with HA under aseptic conditions. For this, the TyrPC NGTs were placed on a mandrel and were subsequently dipped in 1% (w/v) sterile thiol-modified cGMP grade hyaluronan solution (cGMP HyStem, ESI BIO, California, USA) for 30 sec. Next, the NGTs were crosslinked in 1% (w/v) cGMP grade poly(ethylene glycol diacrylate) (PEGDA) solution for 30 sec. The coating was air dried 5 minutes. These steps were repeated 20 times resulting in a ~150–200 μm thick HA coating. After the last coating, NGTs were dipped in sterile cGMP grade hyaluronan solution for 30 sec and then dried overnight in a laminar flow hood. The pore size of the HA coated braided TyrPC NGTs was approximately 2 μm as shown previously [9].

After HA coating, the braided TyrPC NGTs (2.3 mm ID, 1.2 cm long) were sealed in an aluminum pouch and terminally sterilized by electron beam (E-beam) at a radiation dose of 25 KGy (Johnson & Johnson Sterility Assurance (JJSA), Raritan, NJ). The terminally sterilized cGMP HA coated NGTs were tested for sterility using a TSB test. Upon visual observation of the cGMP HA coated NGTs in the TSB test, no turbidity was observed over the 14 days indicating that the cGMP HA coated NGTs were sterile.

### Scanning Electron Microscopy (SEM) Characterization

NGTs were imaged using scanning electron microscopy (SEM, Amray 1830I, 20 kV) after sputter coating with Au/Pd (SCD 004 sputter coater, 30 milliamps for 120 seconds) to assess for any morphological changes. SEM imaging enabled qualitative assessment of the uniformity of the HA coating. For example, if the coating was disrupted, then hole(s) in the coated layer spanning the pores would be observed.

### Operative Technique: 1 cm Segmental Repair of Deep Peroneal Nerve Injury in Swine

All procedures were approved by the University of Pennsylvania’s Institutional Animal Care and Use Committee (IACUC) and adhered to the guidelines set forth in the NIH Public Health Service Policy on Humane Care and Use of Laboratory Animals (2015). This study utilized young-adolescent Yorkshire domestic farm pigs, 3-4 months of age weighing 25–30 kg (Animal Biotech Industries, Danboro, PA). A total of 15 pigs were enrolled to (a) evaluate the mechanism of action across the graft zone at 2 weeks post repair (n=13) and (b) compare nerve conduction and muscle electrophysiological functional recovery at 6 months (n=2). These two time points were selected to demonstrate that nerve repair using TyrPC NGT results in a) no excessive inflammatory responses or implant-associated toxicity at 2 weeks and 6 months post repair; b) ongoing pro-regenerative Schwann cell infiltration and axon regeneration across TyrPC NGTs at 2 weeks post repair; and c) axon maturation and muscle reinnervation at 6 months post repair.

Surgical procedures were performed under general anesthesia. Animals were anesthetized with an intramuscular injection of ketamine (20–30 mg/kg) and midazolam (0.4–0.6 mg/kg) and maintained on 2.0–2.5% inhaled isoflurane/oxygen at 2 L/min. Preoperative glycopyrrolate (0.01–0.02 mg/kg) was administered subcutaneously to control respiratory secretions. All animals were intubated and positioned in left lateral recumbency. An intramuscular injection of meloxicam (0.4 mg/kg) was delivered into the dorsolateral aspect of the gluteal muscle and bupivacaine (1.0–2.0 mg/kg) was administered subcutaneously along the incision site(s) for intra- and post-operative pain management. The surgical site was draped and cleaned under sterile conditions. Heart and respiratory rates, end tidal CO_2_, and temperature were continuously monitored in all animals.

A 10 cm longitudinal incision was made on the lateral aspect of the right hind limb from 1.5 cm distal to the stifle joint, and extending to the lateral malleolus as previously described [14]. In brief, the fascial layer was bluntly dissected and the peroneus longus was retracted to expose the distal aspect of the common peroneal nerve diving into the muscle plane between the extensor digitorum longus and tibialis anterior. Further dissection revealed the bifurcation of the common peroneal nerve (CPN) into the deep and superficial peroneal nerves (DPN). A 1 cm defect was created in the deep peroneal nerve, 0.5 cm distal to the bifurcation. The defect was repaired with either: (1) Neuroflex™ NGT (n=4), made from cross-linked bovine collagen (2 mm diameter x 1.2 cm long; Stryker, Mahwah, NJ), (2) E1001(1k) NGT (n=4; 2 mm diameter x 1.2 cm long), or (3) reverse autograft (n=5). For the NGT repairs, the conduit was secured to the nerve stumps using two 8–0 prolene horizontal mattress sutures, 1 mm in from the end, both proximally and distally. For the reverse autograft repair, the 1 cm segment was removed and rotated so that the proximal nerve stump was sutured to the distal autograft segment and the distal nerve stump was sutured to the proximal autograft segment using two 8–0 prolene simple interrupted sutures on each end [3]. The surgical area was irrigated with sterile saline and the fascia and subcutaneous tissues were closed in layers with 3–0 vicryl interrupted sutures. The skin was closed with 2–0 PDS interrupted, buried sutures, and the area was cleaned and covered with triple antibiotic ointment, a wound bandage, and a transparent waterproof dressing.

In a separate experiment, chronic regeneration and recovery was assessed at 6 months following repair of a 1 cm deep peroneal nerve defect with a sural nerve autograft (n=1) or a E0000 NGT (n=1). In this experiment, E0000 was used as this version is non-degradable, which facilitated localization and visualization of the graft zone at 6 months post repair. The sural nerve was selected as the donor nerve for this study as it is considered the “gold-standard” clinical repair strategy for segmental nerve defects. For the sural nerve autograft harvest, a second longitudinal incision was made approximately 3 cm posterior to the lateral malleolus and parallel to the Achilles tendon. The fascial tissue was dissected to expose the sural nerve running close to the saphenous vein, and a several cm long segment was excised, placed in sterile saline, and then cut into a 1 cm segment for repair of the DPN defect. The deep layers and skin were closed with 3–0 vicryl and 2–0 PDS buried interrupted sutures, respectively.

### Muscle Electrophysiological Evaluation

Electrophysiological evaluation was performed at 6 months post repair of the 1 cm deep peroneal nerve defect repaired with either a sural nerve autograft or a E0000 NGT. Here, the DPN was re-exposed under anesthesia, and the segment containing the repair was carefully freed to minimize tension and isolate it from the surrounding tissues. The nerve was stimulated (biphasic; 1 Hz; 0–1 mA; 0.2 ms pulses) 5 mm proximal to the repair zone with a bipolar electrode with bends fashioned to maintain better contact with the nerve (Medtronic, Jacksonville, FL; #8227410). A ground electrode was inserted into subcutaneous tissue halfway between the electrodes (Medtronic; #8227103). Compound muscle action potential (CMAP) recordings were measured with a bipolar subdermal electrode placed distally in the extensor digitorum brevis muscle belly. The stimulus intensity was increased to obtain a supramaximal CMAP and the stimulus frequency was increased to 30 Hz to achieve maximum tetanic contraction. All CMAP recordings were amplified with 100x gain and recorded with 10–10,000 Hz band pass and 60 Hz notch filters.

Compound nerve action potential (CNAP) recordings were measured 5 mm distal to the repair zone with a bipolar electrode as described above. All CNAP recordings were amplified with 1,000x gain and recorded with 10–10,000 Hz band pass and 60 Hz notch filters.

### Tissue Processing, Histology, and Microscopy

At the terminal time points, animals were deeply anesthetized and transcardially perfused with heparinized saline followed by 10% neutral-buffered formalin using a peristaltic pump. Hind limbs were removed and post-fixed in formalin at 4 °C overnight to minimize handling artifact. The next day, the ipsilateral and contralateral CPN and DPN were isolated and further post-fixed in 10% neutral-buffered formalin at 4 °C overnight followed by 30% sucrose solution until saturation for cryopreservation. Nerves were embedded in optimal cutting temperature embedding media, flash-frozen, and then sectioned longitudinally (20–25 μm) using a cryostat. After rinsing in 1x phosphate buffered saline, nerve sections were blocked and permeabilized at room temperature using 0.3% Triton-X100 plus 4% normal horse serum for 60 minutes. Primary antibodies diluted in phosphate-buffered saline (PBS) + 4% normal horse serum (NHS) were applied to the sections and allowed to incubate at 4°C for 12 hours. After rinsing in 1x PBS, appropriate fluorescent secondary antibodies (Alexa-594 and/or −647; 1:500 in 4% NHS solution) were added at 18–24°C for 2 hours. For sections stained with FluoroMyelin (1:500; ThermoFisher Scientific, Waltham, MA), the solution was applied for 20 minutes at room temperature. The sections were then rinsed in 1x PBS and cover-slipped with mounting medium.

All semi-quantitative measurements were made by one or more skilled technicians who were blinded as to the experimental group being evaluated. To assess acute regeneration at two weeks post repair, host regenerating axons were stained with phosphorylated and unphosphorylated anti-neurofilament-H (SMI-31/32 1:1000, Millipore, Burlington, MA; NE1022/NE1023) and Schwann cells were labeled with anti-S100 (1:500; Dako, Carpinteria, CA; Z0311). Axon regeneration and Schwann cell infiltration was measured by imaging 2–3 serial longitudinal sections at low magnification to find the sections with the most deeply penetrating axons and Schwann cells as measured from the proximal stump for axons, and from the proximal and distal stump for the Schwann cells. The mean rate of acute axon regeneration was then calculated by dividing the mean penetration by the number of days *in vivo* (14). The proportion of Schwann cell infiltration was calculated by adding the proximal and distal penetration and dividing by the total gap length (1 cm) [19]. Axonal maturation and remyelination was evaluated at 6 months with SMI-31/32 and FluoroMyelin, respectively. To evaluate the ECM deposition profiles across the graft region, sections were stained with anti-collagen I (1:500; Abcam, Cambridge, MA; ab34710); anti-collagen III (1:500; Abcam; ab7778); anti-collagen IV (1:500; Abcam; ab6586); anti-laminin (1:500; Abcam; ab11575); and anti-fibronectin (1:500; Abcam; ab6328). The mean intensity and mean intensity relative to the center of the graft were measured for each ECM protein. Several regions of interest (ROI; 100 μm by 100 μm) were from representative locations along the midline or the wall of the graft. Between nine and eighteen representative ROI’s were analyzed and then averaged per animal. To calculate the mean pixel intensity relative to the center of the graft, the average midline intensity (Rm) was subtracted from the averaged edge intensity (Re) and then normalized to the average midline intensity using the following formula: (Re-Rm)/Rm. For instance, a mean normalized value of 0 would represent a homogenous ECM deposition profile between the edge and center of the graft. A mean normalized value of 1 would represent an elevated ECM deposition profile along the edge, whereas a mean normalized value of −1 would represent a decreased ECM deposition along the edge. To assess macrophage localization around the graft region at two weeks following repair, sections were stained with anti-IBA1 (1:1000; Fujifilm Wako, Richmond, VA; 019–19741) and Hoechst 33342 (1:10,000; Invitrogen, Waltham, MA; H3570).

Images were obtained with a Nikon A1R confocal microscope (1024×1024 pixels) with a 10x air objective and 60x oil objective using Nikon NIS-Elements AR 3.1.0 (Nikon Instruments, Tokyo, Japan). To minimize potential bias, a blinded trained researcher performed all subsequent analyses using randomly-coded images separated into individual channels. The mean pixel intensity was measured from images acquired with same confocal settings within regions-of-interest drawn using Nikon NIS-Elements. Mean axon regeneration and Schwan cell infiltration at two weeks were compared using one-way ANOVA followed by Tukey’s multiple comparison test to determine statistical significance (p < 0.05 required for significance; Graphpad, San Diego, CA). Mean intensity and mean intensity relative to the center of the graft for each ECM protein were compared between collagen NGTs and TyrPC NGTs using a two-tailed unpaired Student’s t-test (α = 0.05).

## Results

### Morphometric and Functional Recovery at 6 Months Post Repair

In the 6-month proof-of-concept experiment, nerve electrical conduction and motor functional recovery was demonstrated at 6 months following a 1 cm segmental nerve repair with E0000 and an autograft (**Supplemental Figure 1**). Electrical conduction across the graft and evoked muscle response suggested ongoing reinnervation of the neuromuscular junctions at 6 months for both cases. Histologically, regenerating axons were visualized spanning the entire length of the 1 cm graft with ongoing maturation and myelination at 6 months post repair (**Supplemental Figure 2**). Minimal fibrosis was noted surrounding the repaired nerves, which appeared to be well-vascularized around the graft and more distally. Qualitatively, atrophy of the extensor digitorum brevis muscle appeared consistent between the two animals.

### E1001(1k) NGT Fabrication and Hydrogel Coating Characterization

Electron microscopy (EM) was performed on E1001(1k) NGTs to evaluate the braided micro-structure, as well as to assess the uniformity of the HA hydrogel coating. The HA hydrogel was found to uniformly coat the NGTs (**Figure 1**). E-beam sterilization had no discernible effect on the structure of the NGT or the morphology of HA coating.

**Figure 1.**
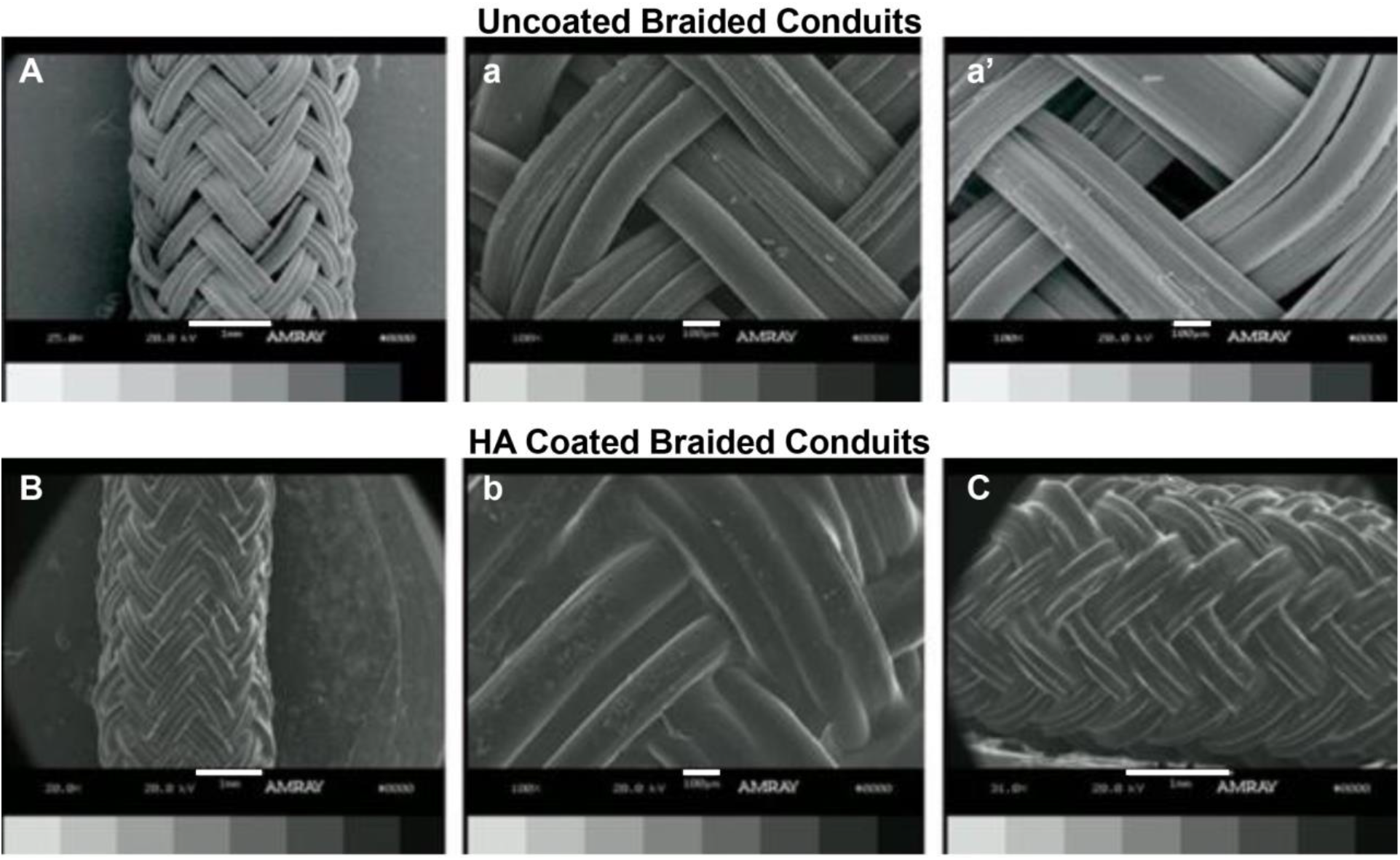
Representative Scanning Electron Microscopy Images of Braided TyrPC NGTs. Scanning electron microscopy (SEM) micrographs of E1001(1k) braided NGTs with triaxial braid shown as uncoated (A) and HA-coated (B). The individual fiber diameter varied from 100 μm–200 μm and the average pore size was 90 μm. Lower magnification SEM revealed the presence of pores spanning the uncoated braided TyrPC NGTs (A), whereas hydrogel filled the open pores in the HA coated braided TyrPC NGTs (B). Representative higher magnification images detailing the open pores (a’, a”) compared to the HA coated pores (b’). The HA coating was uniform and covered the pores completely (B, C). Scale bars: (A, B, C) 1000 μm, (a, a’, b) 100 μm.

### Histological Assessment of Nerve Regeneration at Two-Weeks Post Repair

Initial nerve regeneration into the NGTs was evaluated at two weeks post-repair for Collagen or E1001(1k) NGTs, in comparison to autograft repairs. The main bolus of regenerating axons was visualized across the width of the graft zone of the E1001(1k) NGT, collagen NGT, and autograft repairs. The distance this main bolus of axons had grown at two weeks was measured in the graft region for each group, revealing similar axonal penetration of 3.55 mm ± 0.71 mm for the E1001(1k) NGT, 2.82 mm ± 0.73 mm for the collagen NGT, and 4.32 mm ± 1.29 mm for the autografts, thus yielding statistically equivalent rates of axonal regeneration for all test groups (p=0.24; **Figure 2**). Substantial Schwann cell infiltration was evident throughout the graft zone of all repaired nerves at two weeks, which was not statistically different between the two NGT groups (p=0.18; Figure 3).

**Figure 2.**
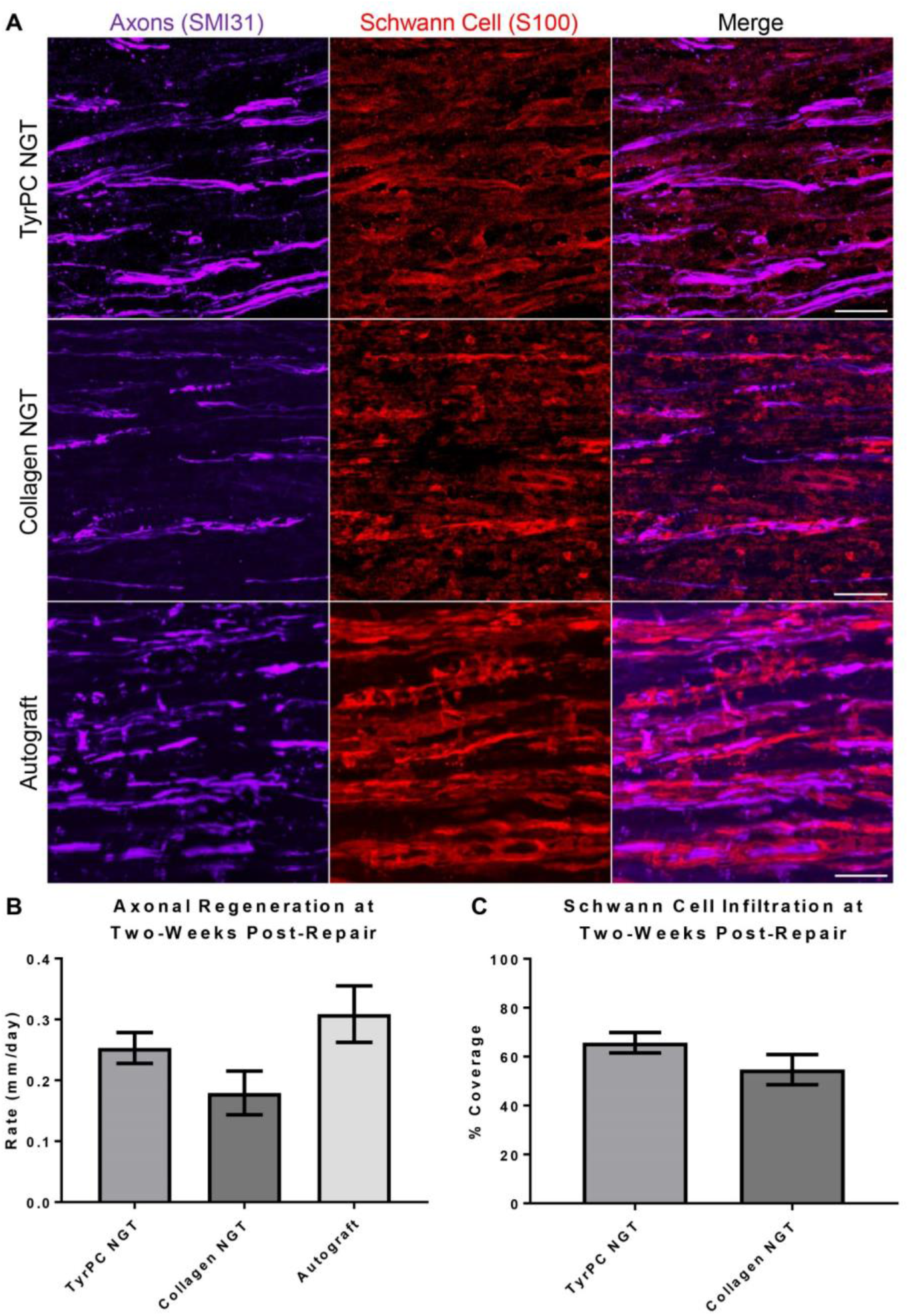
Axon Regeneration and Schwann Cell Infiltration in Nerves Repaired Using Braided E1001(1k) versus Collagen NGTs. (A) At 2 weeks post repair following a 1 cm nerve lesions, longitudinal sections were stained for axons and Schwann cells. (B) Longitudinal axonal outgrowth was measured and rate was calculated by dividing the mean axon penetration by 14 days *in vivo*. (C) Schwann cell infiltration was measured by adding the proximal and distal penetration normalized to the entire length of the graft [19]. Residual Schwann cells were observed throughout the autograft repair, as expected. No significant difference was found between groups for axonal regeneration (p=0.24) or Schwann cell infiltration (p=0.18). Data presented as mean ± SEM. Scale bar: 100 μm.

**Figure 3.**
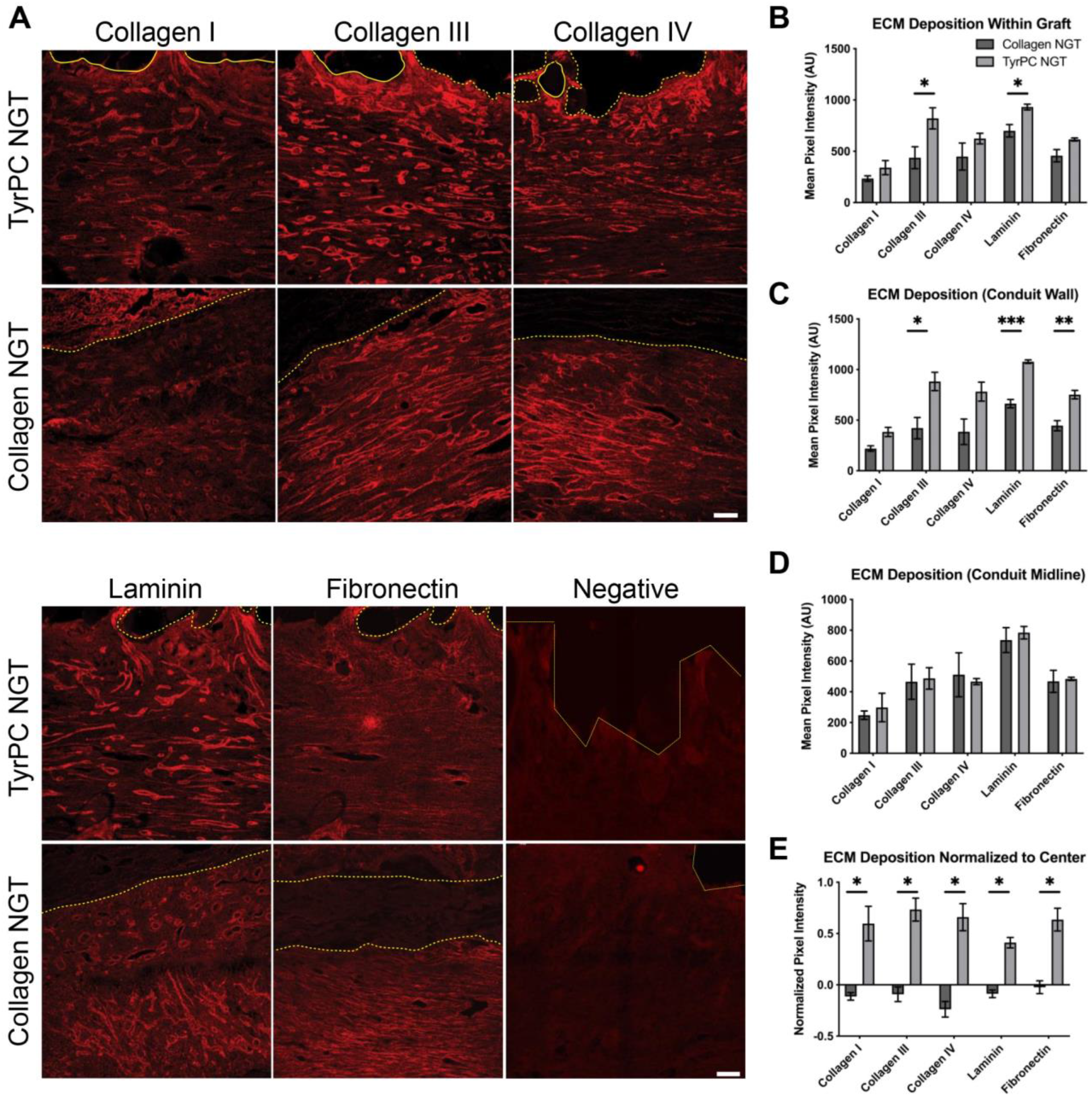
Differential Acute ECM Deposition in Nerves Repaired Using Braided E1001(1k) versus Collagen NGTs. At 2 weeks post repair of 1 cm peroneal nerve lesions, longitudinal sections were labeled for various ECM proteins within braided TyrPC NGTs versus collagen NGTs. (A) Representative images showing ECM protein deposition at the biomaterial interface of the nerve conduits. Dashed lines denote the inner edge of the NGT spanning the graft region. (B-D) Mean pixel intensity of the ECM proteins are shown for different locations within the graft. (B) Within the entire graft region, greater deposition of collagen III and laminin was found in TyrPC NGTs compared to collagen NGTs. (C) At the biomaterial-cell interface, greater deposition of collagen I, collagen III, laminin, and fibronectin were observed in TyrPC NGTs compared to collagen NGTs. (D) No statistical differences in ECM deposition were found at the midline of the conduits. (E) To assess the relative ECM distribution within the conduit, ECM deposition along the wall was normalized to the midline. Greater deposition of collagen I, collagen III, collagen IV, laminin, and fibronectin were found concentrated along the conduit wall relative to the midline region in TyrPC NGTs compared to collagen NGTs. Negative control sections stained with only secondary antibodies are shown for comparison. Statistical comparisons between groups are shown. * p<0.05, ** p<0.01, *** p<0.001. Data presented as mean ± SEM. Scale bar: 100 μm.

After two weeks, various ECM proteins were deposited within the graft region of all NGTs. Differences in the ECM deposition profiles between the groups were apparent both qualitatively and semi-quantitatively (**Figure 3A**). The deposition of specific ECM proteins throughout the entire NGT was analyzed, revealing elevated concentration of collagen III and laminin within E1001(1k) NGTs compared to collagen NGTs (p<0.05 each; **Figure 3B**). Although fibronectin within the entire graft was not statistically significant, there may be a trend towards greater deposition within E1001(1k) NGTs (p=0.08). Next, we investigated whether there were any differences in protein expression localized along the wall at the biomaterial-cell interface and within the midline of the conduit. While there were no statistically significant differences in levels of any of the ECM proteins at the midline of the NGTs (**Figure 3D**), there were greater depositions of collagen I (p<0.05), collagen III (p<0.05), laminin (p<0.001), and fibronectin (p<0.01) along the wall of the E1001(1k) NGTs compared to collagen NGTs (**Figure 3C**). The collagen IV deposition along the wall was not statistically significant but there was a trend towards greater deposition along E1001(1k) NGTs (p=0.07). Notably, when assessing the ECM deposition at the internal NGT surface (at the biomaterial-cell border) relative to the central lumen, we found that levels of collagen I, collagen III, collagen IV, laminin, and fibronectin were particularly elevated in the E1001(1k) NGTs compared to the collagen NGTs (p<0.05 each; **Figure 3E**). Thus, in the case of the collagen NGTs, ECM deposition for these proteins was concentrated in the center of the NGT, away from the walls as indicated by the negative relative pixel intensity values, whereas in the TyrPC case the ECM deposition was concentrated along the walls indicating a potentially broader neurotrophic effect from the material itself. Regenerating axons and infiltrating Schwann cells were found in close proximity to bands of collagen and laminin ECM proteins within the graft zone of both groups (**Figure 4**). In addition, a moderate level of macrophage activity was found in the graft region in both groups. However, qualitatively, there appeared to be greater macrophage localization along the edge of the collagen NGT compared to the edge of the E1001(1k) NGTs (**Figure 5**).

**Figure 4.**
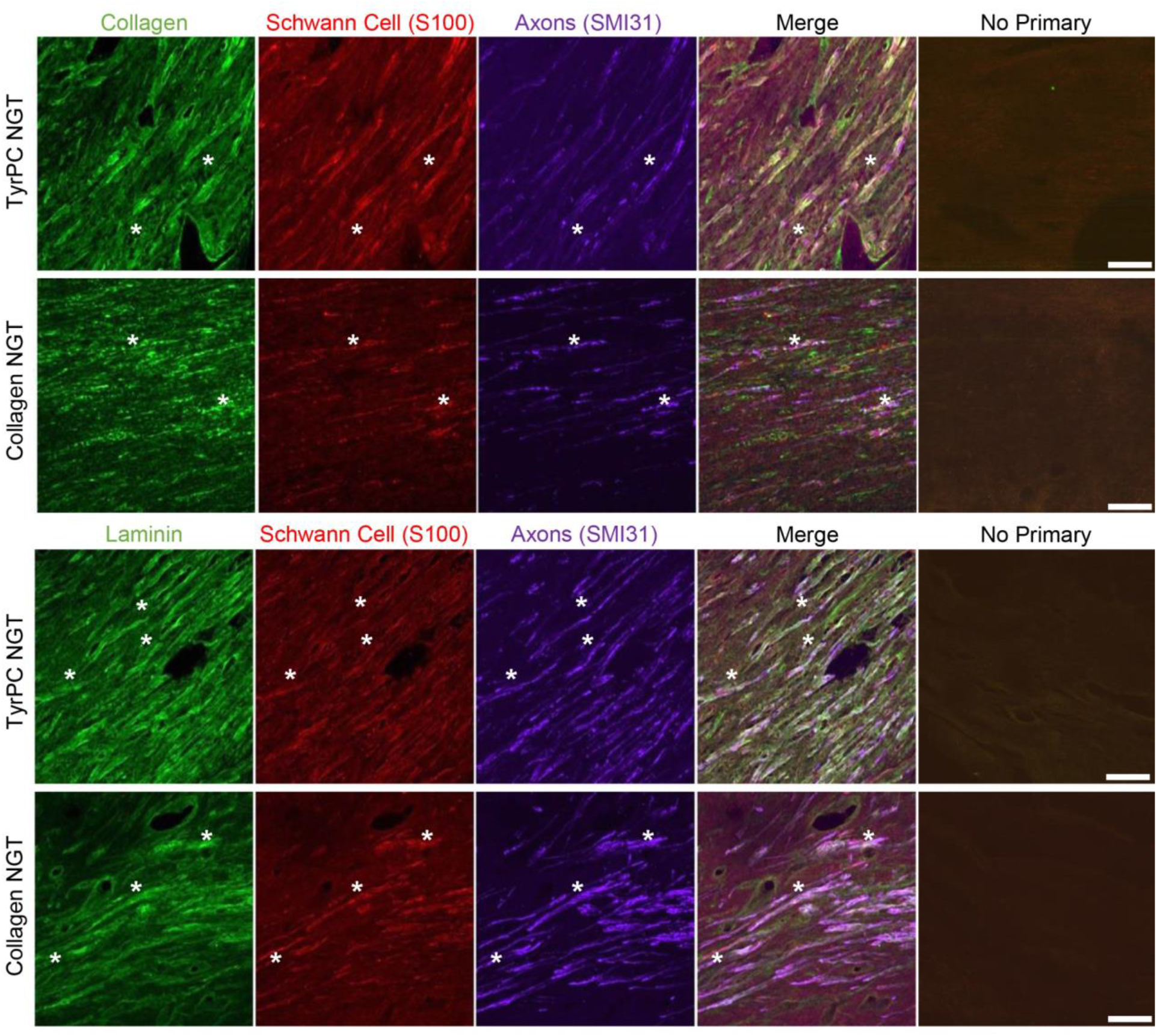
ECM Deposition in Relation to Schwann Cell Infiltration and Axon Regeneration in Nerves Repaired Using Braided E1001(1k) versus Collagen I NGTs. At 2 weeks post repair of 1 cm nerve lesions, representative longitudinal sections showing ECM protein expression and deposition in conjunction with Schwann cell presence and axonal ingrowth. Asterisks denote regions where aligned Schwann cells are in close proximity with regenerating axons. Negative control sections stained with only secondary antibodies are shown for comparison. Scale bar: 100 μm.

**Figure 5.**
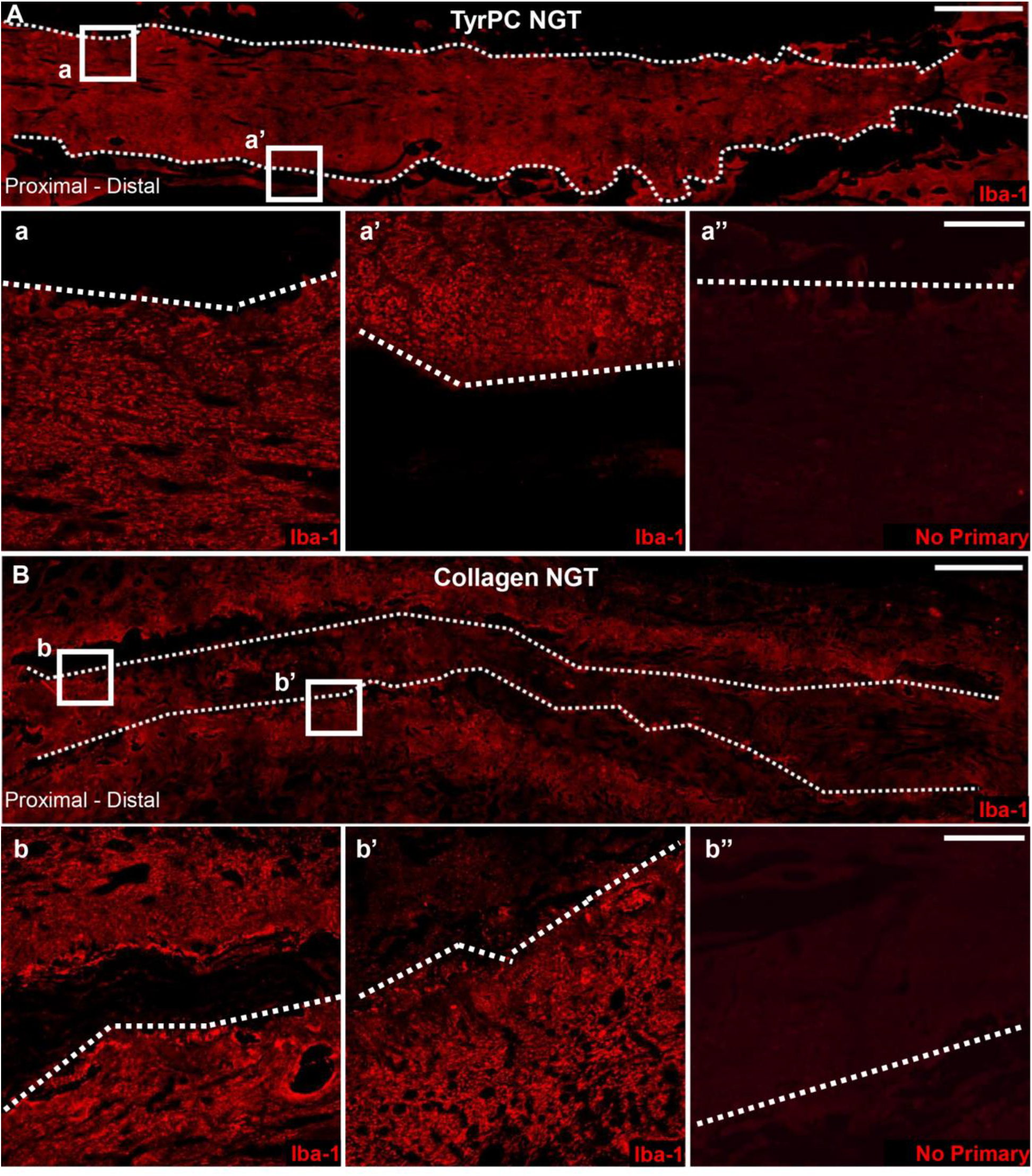
Macrophage Infiltration and Activation Along the Edge of the NGTs Following Nerve Repair Using Braided E1001(1k) versus Collagen NGTs. At 2 weeks post repair, longitudinal nerve sections were stained for macrophages (IBA1) to qualitatively assess macrophage reactivity following repair using (A) TyrPC or (B) collagen NGT. (A-B) Moderate level macrophage activity was visualized along the edge of the NGT in both groups, with a greater build-up of macrophages at the biomaterial-tissue border for the collagen NGT group. Dashed lines denote the inner edge of the NGT spanning the graft region. Negative control sections stained with only secondary antibodies are shown for comparison. Scale bars: (Top, low magnification) 1000 μm, (Bottom, zoom in) 100 μm.

## Discussion

Over the last few decades, several NGTs have been developed as an alternative to the autograft repair strategy. Although multiple commercially available NGTs have been approved for clinical application in the United States, autografts remain the gold standard for repairing nerve gaps longer than 3 cm [20]. While new NGT designs are often evaluated in mouse or other rodent models [13], large animal models are uniquely suitable to replicate clinical scenarios and simulate nerve regeneration in humans. Here, we report the next phase in studying braided TyrPC NGTs for nerve regeneration. In the current study, first we demonstrated TyrPC conduits are suitable for nerve repair in a proof-of-concept chronic porcine study with a non-degradable E0000 NGT.

Electrophysiological functional recordings were observed at 6 months without any noticeable inflammatory responses or implant-associated toxicity. Based on these promising findings, we advanced our previously reported HA coated braided TyrPC NGTs and report similar efficacy with a commercially-available NGTs in a clinically-relevant short gap injury porcine model during the acute regenerative phase. Here, acute regeneration was assessed at two weeks following 1 cm segmental defect repair with a degradable E1001(1k) NGT or commercially-available Stryker Neuroflex NGT made from crosslinked bovine collagen I. Neuroflex was chosen as an experimental benchmark because it is a flexible, semipermeable, resorbable tubular matrix used clinically [21]. We found that both E1001(1k) and collagen NGTs supported Schwann cell infiltration and axon regeneration within the lumen yet prevented deleterious fibrotic invasion at two weeks post repair. Although differences between groups were not found to be statistically significant for these regenerative metrics, E1001(1k) repairs trended towards improved performance.

Recent advancements in bioengineering have allowed for the fabrication of next-generation NGTs with pro-regenerative properties. However, there are several criteria that NGTs must demonstrate prior to clinical deployment, such as appropriate porosity and pore size, mechanical stability, and degradation into resorbable products. Appropriate porosity is necessary to allow for waste and nutrient diffusion into the graft region, however, the pore size should be within the 5–30 μm range to minimize excessive fibrosis and inflammatory cell infiltration [22–24]. Braided biomaterials often provide improved mechanical strength and provide kink-resistance, however, large pores may form allowing for tissue infiltration across the conduit wall into the conduit that can hinder nerve regeneration. To mitigate these concerns, braided TyrPC NGTs were coated with a biocompatible HA-based hydrogel [9, 12]. HA-coated braided TyrPC NGTs exhibit sufficient porosity for mass transport considerations yet have been shown to prevent detrimental cellular infiltration into the graft zone [9]. The HA-coated TyrPC NGT has a pore size of 2 μm [9] whereas the semipermeable collagen NGT has a pore size of 0.1–0.5 μm [25]. Thus, both the HA-coated braided TyrPC and collagen NGTs have a small pore size amenable for oxygen and/or nutrient diffusion, but should prevent unwanted lateral cellular infiltration. In this study, we found the HA coated braided TyrPC NGTs supported axon regeneration and Schwann cell infiltration across the conduit at 2 weeks post repair, suggesting that the HA coating successfully prevented undesirable cellular infiltration across the conduit walls that could have impeded nerve regeneration (e.g., by forming fibrotic tissue within the NGT).

Properly designed NGTs are able to support endogenous nerve regeneration, which proceeds longitudinally across the proximal and distal nerve stumps. Briefly, during the initial regeneration phase following segmental nerve repair using an NGT, fluid rich in pro-regenerative soluble factors/proteins secreted by both nerve stumps begins to rapidly accumulate within the conduit inner lumen [26–28]. Next, a fibrin matrix forms spanning the length of the conduit to provide a substrate for cellular migration. The fibrin scaffold allows for the attachment and longitudinal infiltration of various non-neuronal cells, such as Schwann cells, fibroblasts, macrophages, and endothelial cells, that influence axonal extension and/or maturation within the conduit. A previous study comparing E10–0.5(1k) NGT repair to a non-porous polyethylene NGT in a mouse model has reported differences in the early regenerative phase, including greater fibrin deposition and Schwann cell infiltration, which may have contributed to greater axon regeneration and myelination at the chronic time point [10]. Although we did not directly investigate fibrin deposition, we found no significant differences in axon regeneration or Schwann cell infiltration between E1001(1k) and collagen NGTs. However, future studies will investigate the role of fibrin deposition in porcine nerve regeneration as this mechanism may be a critical component in successful long gap nerve repair [13].

At 2 weeks post repair, ECM deposition within the graft was compared between TyrPC E1001(1k) and collagen NGTs. Over the entire graft region, increased levels of collagen III and laminin were found in the TyrPC NGTs compared to collagen NGTs. Next, we investigated whether there were any spatial differences with the ECM deposition. For all the proteins assessed in this study, the TyrPC NGT and collagen NGT had similar ECM concentrations within the center of the conduit. However, greater levels of collagen III, laminin, and fibronectin concentrations were found in close proximity to the biomaterial-cell layer interface along the wall. From a translational perspective, these findings demonstrate the braided TyrPC NGT did not disrupt the ECM deposition process within the conduit necessary for axon regeneration and Schwann cell infiltration within the graft, yet may passively provide pro-regenerative structural proteins compared to the collagen NGT.

Moreover, by normalizing the ECM deposition along the conduit wall to ECM deposition within the midline, we were able to better contrast the ECM distribution between groups. Indeed, the two NGTs exhibited stark differences in the distribution of ECM proteins within the NGT. Greater concentration of collagen I, collagen III, collagen IV, laminin, and fibronectin were found along the edge of the TyrPC NGT relative to the center. When examined together, these findings suggest there may be enhanced adsorption to the braided TyrPC fibers compared to the cross-linked collagen casing. Our findings corroborate a previous study that demonstrated E10–0.5(1k) 2D films promote selective adsorption of endogenous ECM proteins laminin, fibronectin, and collagen I, and increased Schwann cell process spreading and neurite outgrowth *in vitro* [10]. Collectively, our findings suggest that TyrPC NGTs may create a “biological niche”, or favorable neurotrophic environment, along the inner walls of the NGT that appears to support nerve repair even in the absence of externally added biologics in a clinically-relevant swine model.

Based on our histological findings, regenerating axons do not appear to directly interact with proteins on the conduit walls, yet differences in acute ECM deposition may influence the pro-regenerative environment within the conduit, as suggested by the axon outgrowth and Schwann cell infiltration in the TyrPC NGT trending towards significance. Moreover, differences in pro-regenerative ECM deposition may be a critical early event post implantation that influences longer-term regenerative outcomes by affecting downstream biological events requiring ECM interactions, such as cell attachment and tissue development [29]. Although the exact role is unclear, collagen I/III (fibril forming collagen) is thought to be involved in providing peripheral nerves with mechanical strength and viscoelastic properties [30, 31]. Collagen IV is a major component of the basement membrane and enables Schwann cell attachment and spreading, a critical step in nerve regeneration [30]. Laminin and fibronectin are also ECM proteins that are important structural components of the basement membrane that regulate axon regeneration and influence non-neuronal cells, such as Schwann cells and macrophages during the acute regenerative phase [32]. Laminin can directly modulate Schwann cell proliferation, elongation, and survival, and also promotes neuronal survival *in vitro* [33] and provides axonal guidance [34]. During regeneration, fibronectin is upregulated and increases Schwann cell spread and proliferation, and neurite outgrowth [35, 36]. Besides from providing endogenous support, collagen, laminin, and fibronectin have been thoroughly investigated and shown to improve nerve regeneration in various rodent models as fillers for conduits [37–40], which has led to development of various biomimetic conduits [41].

During the early regenerative phase, conduits provide necessary structural support but then must degrade and resorb to decrease any mechanical mismatch with surrounding soft tissue and/or physical impediments to remodeling. The degradation rate should be tailored such that sufficient regeneration occurs prior to loss of tube mechanical properties, or else loss of NGT patency and corresponding graft failure will ensue [42]. In addition, NGTs should also be flexible and kink-resistant to allow for repair spanning articulating joints, yet must be amenable for suturing to prevent suture pull out and ultimately graft failure. Lastly. NGTs must ultimately degrade into resorbable byproducts, and all materials and degradation products must be biocompatible, non-cytotoxic, and non-immunogenic. The specific chemical composition of TyrPC can be tuned, providing a library of otherwise similar polymers with varying degradation and resorption rates. Braided TyrPC NGTs also offer a very high degree of kink-resistance, allowing for over 120° of flexing without luminal occlusion [9]. Earlier small animal studies have demonstrated that TyrPC NGTs significantly enhance regeneration and functional recovery, and exhibit excellent mechanical strength, suturability, and flexibility [9–12].

Macrophages play an important role during nerve regeneration by clearing debris and remodeling ECM to facilitates Schwann cell infiltration into the graft region [43]. In this study, macrophages were more concentrated around the edge of the collagen NGT, whereas the distribution of the macrophages appeared more diffuse in the E1001(1k) NGT. The presence of macrophage infiltration along the edges of the NGTs appeared consistent with other studies that have demonstrated NGTs can illicit an immunological response following repair [44]. Moreover, future work is necessary to determine if the biomaterial may elicit differential host macrophage responses, such as fully encapsulating the NGT, propagating inflammatory cascades, and/or facilitating angiogenesis and axon regeneration. Here, investigations exploring different balances along the continuum of M1 and M2 macrophage phenotypes within the graft and along the edges of the E1001(1K) and collagen NGTs would be warranted. The implications of these findings and whether there is a relationship with the differences in cell-material interactions and ECM deposition are currently unclear, but should be established in future studies.

In this study, we demonstrated that TyrPC NGTs may be a suitable alternative to commercially-available NGTs in a clinically-relevant short gap injury porcine model. To date, there are no commercially-available NGTs indicated for nerve repair greater than 3 cm. A major limitation for long gap nerve repair using NGTs is due to the slow axonal regeneration and Schwann cell infiltration within the graft. Although both TyrPC NGTs and collagen NGTs demonstrated similar regeneration during the acute repair phase, the increased adsorption of the pro-regenerative ECM proteins along the wall of the TyrPC NGT may enable greater functional recovery at later time points. In addition to the intrinsic challenges associated with axon regeneration, physical considerations, such as nutrient diffusion, mechanical mismatch, and degradation, become increasingly important in clinically-relevant long gap nerve injury. Although for short gap nerve repairs, nutrients can reach the center of the graft via longitudinal diffusion from the terminal ends of the conduit, long gap nerve repair requires mass transport of nutrients across the conduit wall to sustain axon and tissue regeneration along the length of the graft [9, 45]. Therefore, compared to the crosslinked collagen NGT, the slightly larger pore size in the wall of the braided HA-coated TyrPC NGTs may be beneficial for long gap nerve repair. Braided TyrPC NGTs also provide greater kink-resistance than the collagen NGT, which would provide greater mechanical stability for long gap repairs spanning articulating joints. Moreover, further refinements and/or modifications, such as cell supplementation or growth factor delivery, may enhance the ability of TyrPC NGTs as a suitable strategy for long gap nerve repair. Therefore, future work is necessary to compare the regenerative efficacy of braided TyrPC NGTs at chronic time points and in longer gap nerve injuries to less permeable commercially-available NGTs, and to investigate whether differences in ECM protein deposition within the conduit confer any advantages for these long gap repair scenarios.

## Conclusions

Our data suggests that braided TyrPC NGTs coated with crosslinked HA are biocompatible in large mammals, facilitate nerve regeneration comparable to commercially-available NGTs, and prevent deleterious cellular infiltration while likely allowing nutrient exchange. In our 6-month chronic proof-of-concept study, non-degradable E0000 TyrPC and autograft had qualitatively similar levels of functional recovery. In addition, robust nerve regeneration was observed within the braided E1001(1k) TyrPC NGTs at two weeks post repair. To our knowledge, these data represent the first report of a fully synthetic braided NGT that performed similarly and trending towards outperforming an NGT fabricated from collagen, a biologically-sourced material with known neural cell-supportive bioactivity, including promotion of neurite outgrowth. While the exact mechanism underlying the performance of TyrPC NGTs remains unclear, differences in ECM deposition compared to commercially-available collagen NGTs may influences Schwann cell infiltration, axon regeneration, maturation, and ultimately functional recovery. These studies broadly suggest that NGTs fabricated from members of the TyrPC library elicit pro-regenerative host responses and promote functional recovery following short-gap nerve injury. Furthermore, synthetic materials, such as TyrPCs, offer advantages over biologics, including reduced batch-to-batch variability and potentially reduced cost following commercial scale-up. Future studies are warranted to thoroughly assess the regenerative efficacy of braided TyrPC NGTs at chronic time points and in longer gap nerve injuries, and to probe potential molecular mechanisms underlying enhanced ECM protein deposition at the material/host interface. In conclusion, TyrPCs are promising materials for fabrication of NGTs with significant translational potential.

## Acknowledgements

Financial support was provided by the U.S. Department of Defense [CDMRP/JPC8-CRMRP W81XWH-16-1-0796 (Cullen), MRMC W81XWH-15-1-0466 (Cullen), JWMRP W81XWH-14-1-0100 (Kohn & Cullen), AFIRM W81XWH-08-2-0034 (Kohn & Cullen)]. Additional support was provided by RESBIO - The National Resource for Polymeric Biomaterials funded by the National Institutes of Health (EB001046) (Kohn).

## Author Contributions

D.K.C., J.K., and H.K. conceived of and designed experiments. D.B., S.M., H.K., and J.K. designed and carried out TyrPC NGT fabrication and *in vitro* assessment. Z.A., J.R., and H.K., assisted with initial surgical implementation and/or electrophysiological methodology. J.C.B., J.D., and K.D.B. performed nerve repair surgeries and electrophysiology assessments. J.C.B., D.P.B., and F.A.L., performed histological assessment and confocal imaging, and figure preparation. J.C.B. and D.P.B. analyzed data. J.C.B. and D.K.C. conducted initial figure and manuscript preparation. D.K.C. and J.K. oversaw all studies and manuscript preparation. All authors provided edits and comments to the manuscript.

## Competing Financial Interests

The authors confirm that there are no known conflicts of interest associated with this publication and there has been no significant financial support for this work that could have influenced its outcome.

## Data Availability

The raw/processed data required to reproduce these findings cannot be shared at this time as the data also forms part of an ongoing study. Requests for raw/processed data may be addressed to the corresponding author.

---

**Supplemental Figure 1.**
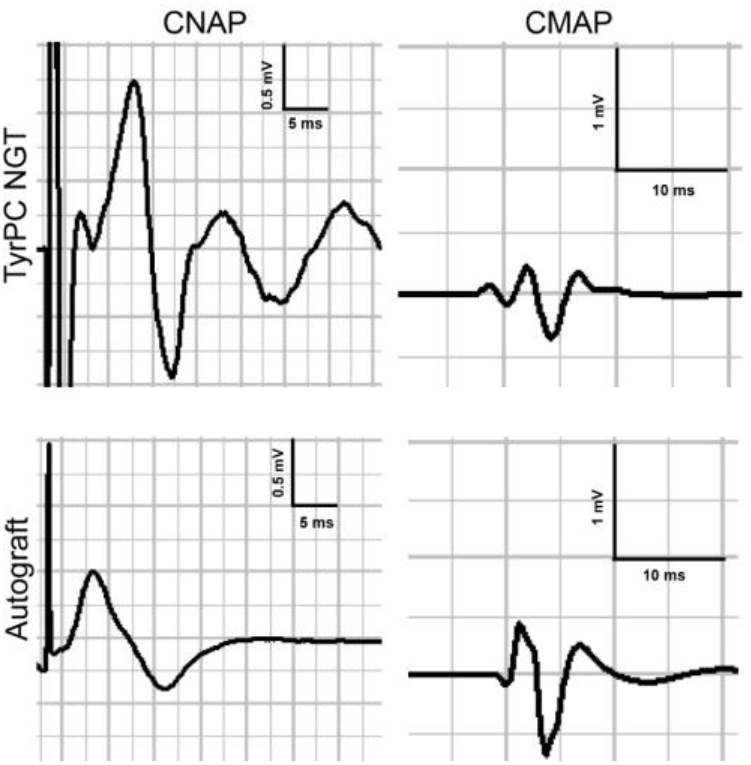
Functional Recovery at 6 Months Post Repair of a 1 cm defect with E0000 NGT and Autograft. Nerve lesions 1 cm in length were repaired with either (A) E0000 NGT or (B) reverse autograft or (1.2 cm long; HA-coated). Functional recovery was assessed at 6 months post repair. Nerve conduction (CNAP) and muscle reinnervation (CMAP) recordings were measured from nerves repaired with either E0000 or autograft repairs. At 6 months post repair, the robust CNAP response showed successful electrophysiological conduction across the graft with regenerating axons undergoing maturation, which was corroborated by the histological findings. Moreover, the evoked muscle response indicated successful reinnervation of neuromuscular junctions in the distal muscle end target at 6 months.

**Supplemental Figure 2.**
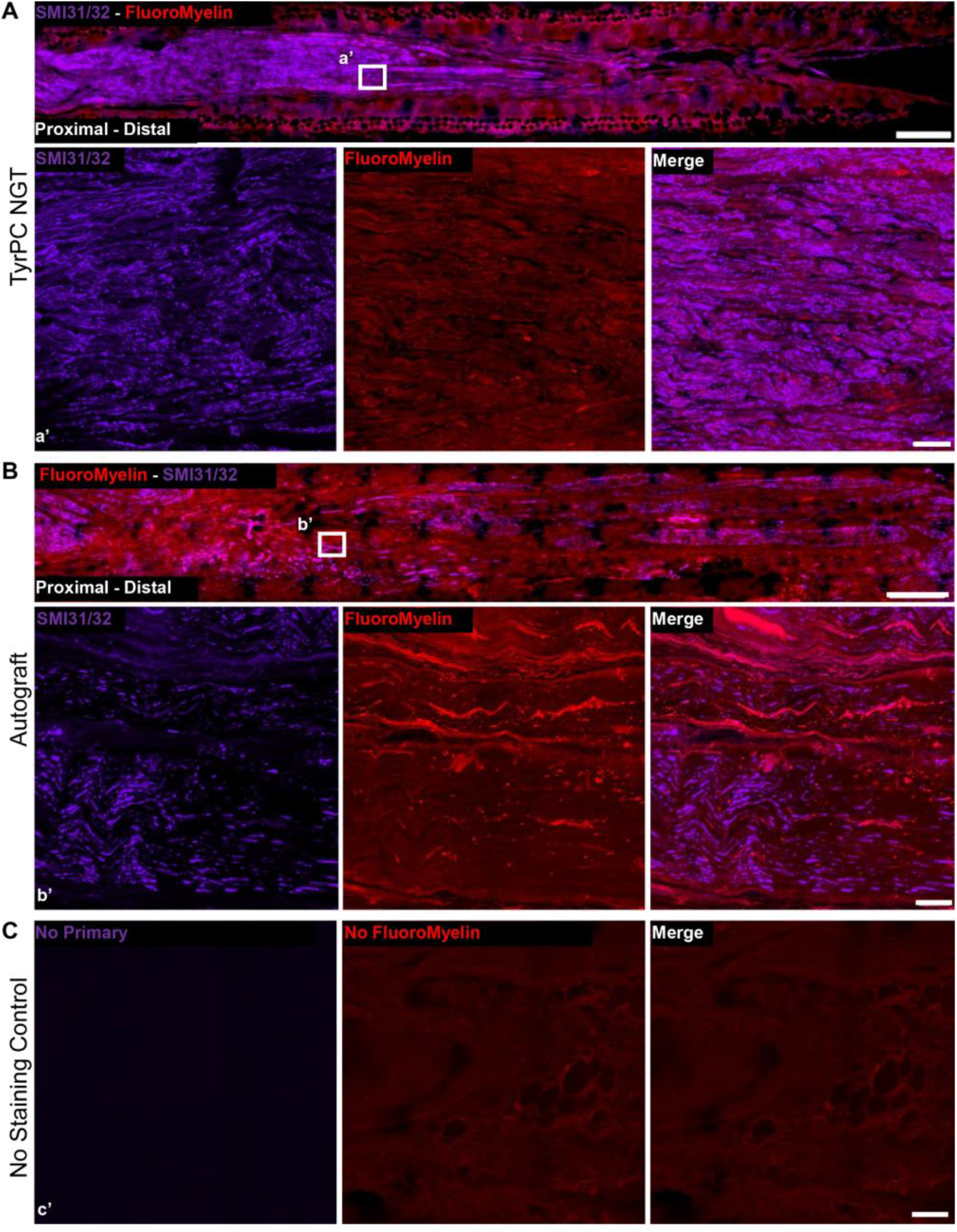
Nerve Regeneration and Myelination at 6 Months Post Repair of a 1 cm defect with E0000 NGT and Autograft. Nerve lesions 1 cm in length were repaired with either (A) E0000 NGT or (B) reverse autograft or (1.2 cm long; HA-coated). Nerve morphometry was assessed at 6 months post repair. Robust axonal regeneration and remyelination were found within both graft regions by labeling for SMI-31/32 and FluoroMyelin, respectively. Regenerating axons were found spanning the entire length of both 1 cm grafts at 6 months post repair. Evidence of the nondegradable TyrPC NGT can be seen in red autofluorescence. (C) TyrPC NGT staining control sections that lacked SMI31/32 application but were incubated with the secondary antibody. FluoroMyelin, which is a fluorescent dye, does not require a secondary antibody for visualization, and therefore it was simply not applied in the staining control section. Scale bars: (Top, low magnification) 1000 μm, (Bottom, zoom in) 100 μm.

## Notes

### Competing Interest Statement

The authors have declared no competing interest.

